# Gap junction protein INNEXIN2 modulates the period of free-running rhythms in *Drosophila melanogaster*

**DOI:** 10.1101/2020.04.29.065037

**Authors:** Aishwarya Ramakrishnan, Vasu Sheeba

## Abstract

The circadian neuronal circuit of *Drosophila melanogaster* is made up of about 150 neurons, distributed bilaterally and distinguished into 7 clusters. Multiple lines of evidence suggest that coherent rhythms in behaviour are brought about when these clusters function as a network. Although chemical modes of communication amongst circadian neurons have been well-studied, there has been no report of communication via electrical synapses made up of gap junctions. Here, we report for the first time that gap junction proteins – Innexins play crucial roles in determining the period of free-running activity rhythms in flies. Our experiments reveal the presence of gap junction protein INNEXIN2 in the ventral lateral neurons. RNA-interference based knockdown of its expression in circadian pacemakers slows down the speed of locomotor activity rhythm. Concomitantly, we find alterations in the oscillation of a core-clock protein PERIOD and in the output molecule Pigment Dispersing Factor in the circadian pacemaker neuron network.

## Introduction

*Drosophila* has been widely used as a model organism in circadian biology because of its robust and easily quantifiable behaviours and relatively few number of neurons controlling them (Allada & Chung, 2010). The adult *Drosophila* circadian circuit is composed of about 150 neurons distributed bilaterally in the brain. Based on their location, they can be divided into lateral neurons (LN) and dorsal neurons (DN). The lateral neurons are further divided into the small ventral lateral neurons (s-LNv), large ventral lateral neurons (l-LNv), the lateral dorsal neurons (LNd) and the lateral posterior neurons (LPN). The dorsal cluster of neurons are further divided into 3 groups as dorsal neurons 1-3 (DN1-3) (reviewed in Sheeba, 2008).

Each of these neurons have a ticking molecular clock composed of a self-sustained transcriptional translational feedback loop (TTFL) made up of four core clock genes *Clock, Cycle, Period* and *Timeless*. The period of these molecular oscillations in mRNA and protein within the pacemaker circuit in the fly brain mirror the period of rhythmic activity-rest behaviour reviewed in (Hardin, 2005). Although molecular circadian clocks in individual neurons can be thought of as ticking cell autonomously due to the precisely timed cycling of their mRNA and proteins, one interesting question that remains to be fully understood is how these distinct neuronal clusters, with distinct intrinsic periodicities (Yoshii et al., 2009) together bring about one coherent period of the behavioural activity rhythm. Early studies of *Drosophila* clock neuronal network have shown that under constant darkness and constant temperature (DD 25 °C), s-LNv neurons and clocks in these cells are necessary and sufficient for the persistence of activity-rest rhythms (Helfrich-Förster, 1998, Renn et al., 1999). s-LNv release neuropeptide Pigment Dispersing factor (PDF) in the dorsal part of the brain via their projections in a time-of-day dependent manner (Park et al., 2000). Lack of PDF results in arrhythmicity of activity-rest rhythms under constant conditions (Renn et al., 1999) suggesting that PDF is necessary for persistence of rhythms. PDF receptor (*PdfR*) is widely distributed in most clock neurons in the circuit (Hyun et al., 2005, Mertens et al., 2005), most of them being responsive to PDF (Shafer et al., 2008), thus establishing its role as an important synchronizing factor in the circadian circuit. Apart from PDF, several other neuropeptides and neurotransmitters have been implicated to play diverse roles in communication among circadian neurons, although none of them have been shown to play as prominent roles as PDF (reviewed in Beckwith & Ceriani, 2015).

Across organisms, and behaviours, most studies have focused on the role played by chemical synapses among neurons in a circuit, even though electrical and chemical synapses have been known to co-exist in neural networks of most organisms (reviewed in Pereda, 2014, Nagy et al., 2018). Gap junctions are tightly coupled clusters of proteins which form intercellular membrane spanning channels connecting the cytoplasm of adjacent cells. These channels facilitate electrical coupling of adjacent cells through diffusion of ions, metabolites and cyclic nucleotides (Faber & Pereda, 2018). Three gene families are known to form gap-junction channels. *Connexins* form gap junctions in chordates; *Innexins* were identified as the structural proteins of gap junctions in invertebrates and *Pannexins* which are structurally similar to *Innexins* are found in some invertebrates and chordates and mostly function as hemichannels (Beyer & Berthoud, 2017). Structurally, gap junction proteins are four-pass transmembrane (TM) proteins with intracellular N and C termini, two extracellular loops and one intracellular loop. *Drosophila melanogaster* has eight members of the *Innexin* family named *Innexin1-8* (reviewed in Bauer et al., 2005). Gap junction hemichannels can be classified as homomeric or heteromeric, composed of the same or different classes of *Innexins* respectively. Intercellular channels are called homotypic if each of the two hemichannels are made of same type of *Innexins* and heterotypic if made of two different hemichannels. The proper physiological functioning of *Innexin* channels is dependent on their specific combinations (Stebbings et al., 2000). Functions of several of the *Innexin* classes have been characterized extensively during development and recently in behaviours exhibited by adult flies (reviewed in Güiza et al., 2018, summarized in Table.1).

Most of the studies which suggest a role of gap junctions in modulating circadian behaviour have been undertaken in mammals. Neurons in Suprachiasmatic nucleus (SCN); the circadian pacemaker in mammals are well-synchronized and form a coherent oscillator network (Herzog et al., 2017). Even though various mechanisms for intercellular coupling of SCN neurons have been suggested, including GABA and Vasoactive Intestinal peptide (VIP) (reviewed in Evans & Gorman, 2016), circadian clocks in the developing SCN are fully functional and apparently synchronized long before neurochemical synaptic connections are present (Moore & Bernstein, 1989, Landgraf et al., 2014), suggesting the existence of an additional mechanism for cell communication within the SCN like gap junctions. The SCN expresses a number of different *Connexins* (Welsh & Reppert, 1996, Colwell, 2000, Rash et al., 2007). Gap junction coupling in intact SCN tissue has been demonstrated with tracer molecules and by electrical stimulation and recording of neighbouring cells,(Jiang et al., 1997, Colwell, 2000, Shinohara et al., 2000) Gap junctions are involved in synchronous firing of coupled cells in SCN neuronal network which can be suppressed with Carbenoxolone, a reversible blocker of gap junctions (Wang et al., 2014). In particular, *Connexin*36 (*Cx36*) has been reported to play a crucial role in electrical coupling of SCN neurons. Knockout of *Cx36* blocks intercellular electrical coupling between SCN neurons, and adult *Cx36* knockout mice display a lower amplitude of circadian locomotor activity rhythms and a decrease in overall activity levels under constant environmental conditions (Long et al., 2005). Another recent study, however shows that absence of *Cx36* does not affect the synchronous PER protein oscillations in SCN neuronal network even though the *Cx36* knockout mice have lengthened period of wheel-running activity as compared to controls (Diemer et al., 2017). Thus, even though some evidences suggest the importance of electrical coupling among SCN neurons for synchronous firing as well as for output behaviour, the underlying mechanisms are poorly understood. To the best of our knowledge, in invertebrates, there has been only one study in the cockroach species, *Leucophaea maderae*, where, use of gap junction blockers results in desynchronized firing of accessory medulla neurons which are the circadian pacemaker centre (Schneider & Stengl, 2006). However, despite nearly three decades of studies of the *D. melanogaster* circadian pacemaker circuit, there has been no report of involvement of gap junctions in modulating circadian rhythms.

We performed a screen to identify the role played by gap junction proteins in the *Drosophila* circadian circuit. We found that the levels of two gap junction genes *Innexin1* and *Innexin2* determine a very core clock property, the free-running period. Here we have report results which demonstrate that INNEXIN2 is expressed in circadian pacemaker neurons and its knockdown causes activity rhythm to slow down. We report that oscillation of the core molecular clock protein (PERIOD) is lengthened in most circadian neurons upon *Innexin2* knockdown along with a change in the levels of neuropeptide PDF in the s-LNv dorsal projections. Thus, we provide the first evidence of a role for gap junction proteins in circadian pacemaker circuit of *Drosophila melanogaster* and suggest possible mechanisms for the same.

## Materials and Methods

### Fly lines

All genotypes were reared on standard cornmeal medium under LD (12 hr **L**ight: 12 hr **D**ark) cycles and 25 °C. The following fly lines were used in this study; *w*^*1118*^ (BL 5905), UAS *Innexin1* RNAi (BL 44048), UAS *Innexin2* RNAi (BL 42645), UAS *Innexin3* RNAi (BL 60112), UAS *Innexin4* RNAi (BL 27674), UAS *Innexin5* RNAi (BL 28042), UAS *Innexin6* RNAi (BL 44663), UAS *Innexin7* RNAi (BL 26297), UAS *Innexin8* RNAi (BL 57706), UAS *GFP-NLS* (BL 4776*), tub GAL80*^*ts*^ (BL 7017), *Clk4.1M GAL4* (BL 36316), *Clk4.5F GAL4* (BL 37526), *repo GAL4* (BL 7415) (Bloomington *Drosophila* Stock Centre, Indiana). *Pdf GAL4* and *tim* (A3) *GAL4* (obtained from Todd Holmes, UC Irvine), *Clk856 GAL4* (provided by Orie Shafer, ASRC, CUNY), *pdf GAL80* (obtained from Helfrich-Forster, University of Wurzburg), *dvpdf GAL4* (obtained from Michael Rosbash, Brandeis University).

### Locomotor activity rhythm assay

Individual virgin male flies (4-6 days old) were housed in glass tubes (length 65mm, diameter 7mm) with corn food on one end and cotton plug on the other end. Locomotor activity was recorded using the ***D****rosophila* **A**ctivity **M**onitors (DAM, Trikinetics, Waltham, United States of America). Experiments were conducted in incubators manufactured by Sanyo (Japan) or Percival (USA).

### Activity data analysis

Raw data obtained from the DAM system were scanned and binned into activity counts of 15 minute intervals. Data was analysed using the CLOCKLAB software (Actimetrics, Wilmette, IL) or RhythmicAlly (Abhilash & Sheeba, 2019). Values of period and power of rhythm were calculated for a period of 7-10 days using the Chi-square periodogram with a cut-off of *p=0.05*. The period and power values of all the flies for a particular experimental genotype were compared against the parental controls using one-way ANOVA with genotype as the fixed factor followed by post-hoc analysis using Tukey’s Honest Significant Difference (HSD) test.

### Immunohistochemistry

Adult *Drosophila* brains were dissected in ice-cold Phosphate buffered saline (PBS) and fixed immediately after dissection in 4% Paraformaldehyde (PFA) for 30 min. The fixed brains were then treated with blocking solution (10% horse serum) for 1-h at room temperature and additional 6-h at 4 °C (Additional incubation is only given in case of staining with anti-PER antibody to reduce staining of non-specific background elements), followed by incubation with primary antibodies at 4 °C for 24-48 h. The primary antibodies used were anti-PER (rabbit, 1:20,000, kind gift from Jeffrey Hall, Brandeis University), anti-PDF (mouse, 1:5000, C7, DSHB), anti-GFP (chicken, 1:2000, Invitrogen), anti-INNEXIN2 (guinea pig, 1:50, kind gift from Michael Hoch, University of Bonn). After incubation, the brains were given 6-7 washes with 0.5% PBS + Triton-X (PBT) after which they were incubated with Alexa-fluor conjugated secondary antibodies for 24-h at 4 °C. The following secondary antibodies were used, goat anti-rabbit 488 (1:3000, Invitrogen), goat anti-mouse 546 (1:3000, Invitrogen), goat anti-mouse 647 (1:3000, Invitrogen), goat anti-guinea pig 546 (1:3000, Invitrogen). The brains were further washed 6-7 times with 0.5% PBT and cleaned and mounted on a clean, glass slide in mounting media (7:3 glycerol: PBS). Exact same procedure was followed for experiments where immunostaining of larval brains (L3 stage) was required.

### Image acquisition and analysis

The slides prepared for immunohistochemistry were imaged using confocal microscopy in a Zeiss LSM880 microscope with 20X, 40X (oil-immersion) or 63X (oil-immersion) objectives. Image analysis was performed using Fiji software (Schindelin et al., 2012). In the samples, clock neurons were classified based on their anatomical locations. PER intensity in these neurons were measured by selecting the slice of the Z-stack which shows maximum intensity, drawing a Region of Interest (ROI) around the cells and measuring their intensities. 3-6 separate background values were also measured around each cell and the final intensity was taken as the difference between the cell intensity and the background. Similar procedure was followed for measuring the intensity of PDF in s-LNv dorsal projections. The intensity values obtained from both the hemispheres for each cell type for each brain was averaged and used for statistical analysis. We used a COSINOR based curve-fitting method (Cornelissen, 2014) to estimate different aspects of rhythmicity like presence of a 24-h periodicity and phase and amplitude values of the oscillation. COSINOR analysis was implemented using the CATCosinor function from the CATkit package written for R (Lee Gierke & Cornelissen, 2016).

## Results and Discussion

### RNAi knockdown screen of *Innexins* in clock neurons

To examine the role of *Innexin*s in the fly circadian network, we performed a RNAi knockdown screen where we knocked down the expression of each of the eight classes of *Innexin* genes using a broad *GAL4* driver that targets all the 150 clock neurons and examined rhythm properties under constant darkness (DD 25°C) (Table 2). In each case, knockdown of *Innexin1* (BL 44048) or *Innexin2* (BL 42645) with *timA3 GAL4*, we observed a significant lengthening of free-running period as compared to its parental controls (Fig. 1A, top; Fig. 1B, left) suggesting that *Innexin1* and *Innexin2* play important roles in determining the period of free-running rhythm. We also quantified the power of the rhythm, which is indicative of the robustness of the underlying clock, (reviewed in Klarsfeld et al., 2003). We found a significant decrease in the power of the rhythm in case of knockdown of *Innexin*7 in clock neurons in one trial (Fig. 1A, bottom). However, this result was not consistent across multiple replicate experiments, and hence it was not pursued further. Importantly, we found no difference in the power of the rhythm in case of *Innexin1* and *Innexin2* knockdown although there was a significant lengthening of period (Fig. 1A, bottom; Fig. 1B, right) suggestive of the fact that the robustness of the clock was not affected. In this report, we describe our studies on the role of *Innexin2* in the circadian clock network. (Studies on the role of *Innexin1* in the circadian network are ongoing and will be described in detail elsewhere, Ramakrishnan and Sheeba, *manuscript in preparation*).

**Table 1:** Known roles of *Innexins* in flies. Summary of the different functional roles played by all eight *Innexin* genes in *Drosophila melanogaster* both during development and adult stages.

| Gene | Functions | References |
| --- | --- | --- |
| *Innexin1*<br>(*Ogre*) | Development of nervous system and optic lobes | Lipshitz & Kankel, 1985<br>Holcroft et al., 2013 |
|  | Escape response to sound stimuli | Pézier et al., 2016 |
| *Innexin2* | Development of gut, tracheal systems, salivary glands, epithelial tissue morphogenesis | (Bauer et al., 2003)<br>(Bauer et al., 2004) |
|  | Development of nervous system | (Holcroft et al., 2013) |
|  | Development of eye | Richard & Hoch, 2015 |
| *Innexin3* | Dorsal closure in embryonic stages | (Giuliani et al., 2013) |
|  | Escape response to sound stimuli | Pézier et al., 2016 |
| *Innexin4*<br>(*Zpg*) | Germ line cell development | Bohrmann &<br>Zimmermann, 2008 |
| *Innexin5* | Visual learning and memory | Liu et al., 2016 |
| *Innexin6* | Associative learning and anaesthesia-resistant memory | (C. L. Wu et al., 2011) |
|  | Visual learning and memory | (Liu et al., 2016) |
|  | Escape response to sound stimuli | (Pézier et al., 2016) |
|  | Modulating behavioural responses during sleep | Troup et al., 2018 |
| *Innexin7* | Development of nervous system | (Ostrowski et al., 2008) |
|  | Associative learning and anaesthesia-resistant memory | C. L. Wu et al., 2011 |
| *Innexin8*<br>(*ShakB*) | Synaptic transmission in the giant fibre system (GFS) | (Phelan et al., 2008) |
|  | Transmission of olfactory information in the antennal lobe | Yaksi & Wilson, 2010 |

**Table 2:** Effect of knockdown of *Innexins* in circadian pacemakers using *tim GAL4* d*river* on free running rhythm properties. Table representing the average period (±SEM), power of the periodogram (± SEM) and % rhythmicity values of all the experimental (*tim; dcr > Inx1-8 RNAi*) lines used for the screen and their respective parental controls (*w; tim GAL4; dcr*) and UAS control (UAS *Inx 1-8 RNAi*).

| Genotype | Period | Power of rhythm | % Rhythmicity |
| --- | --- | --- | --- |
| *w; tim GAL4; dcr* | 23.65 ± 0.01 | 419.2 ± 13.19 | 100 |
| *tim; dcr > Inx1 RNAi* | 25.78 ± 0.07 | 385.3 ± 14.55 | 100 |
| *tim; dcr > Inx2 RNAi* | 24.61 ± 0.07 | 377 ± 13.5 | 100 |
| *tim; dcr > Inx3 RNAi* | 23.62 ± 0.01 | 409.4 ± 16.76 | 100 |
| *tim; dcr > Inx4 RNAi* | 23.87 ± 0.09 | 325.8 ± 14.28 | 81.8 |
| *tim; dcr > Inx5 RNAi* | 23.52 ± 0.03 | 321.2 ± 6.9 | 95.23 |
| *tim; dcr > Inx6 RNAi* | 23.8 ± 0.06 | 376.5 ± 18.17 | 95.23 |
| *tim; dcr > Inx7 RNAi* | 23.73 ± 0.05 | 337.2 ± 91.18 | 100 |
| *tim; dcr > Inx8 RNAi* | 23.58 ± 0.04 | 461.3 ± 13.44 | 100 |
| *yscv; +; Inx1 RNAi* | 23.5 ± 0.03 | 393.8 ± 13.24 | 100 |
| *yscv; +; Inx2 RNAi* | 23.54 ± 0.02 | 424.2 ± 16.55 | 92.59 |
| *yscv; +; Inx3 RNAi* | 23.56 ± 0.02 | 466.1 ± 20.9 | 100 |
| *yscv; +; Inx4 RNAi* | 23.49 ± 0.03 | 327.5 ± 11.34 | 95.83 |
| *yscv; +; Inx5 RNAi* | 23.19 ± 0.04 | 314.6 ± 14.84 | 96 |
| *yscv; +; Inx6 RNAi* | 23.43 ± 0.02 | 437.2 ± 14.4 | 100 |
| *yscv; +; Inx7 RNAi* | 23.56 ± 0.03 | 410.9 ± 51.2 | 96.15 |
| *yscv; +; Inx8 RNAi* | 23.5 ± 0.03 | 454.6 ± 19.17 | 100 |

**Figure 1:**
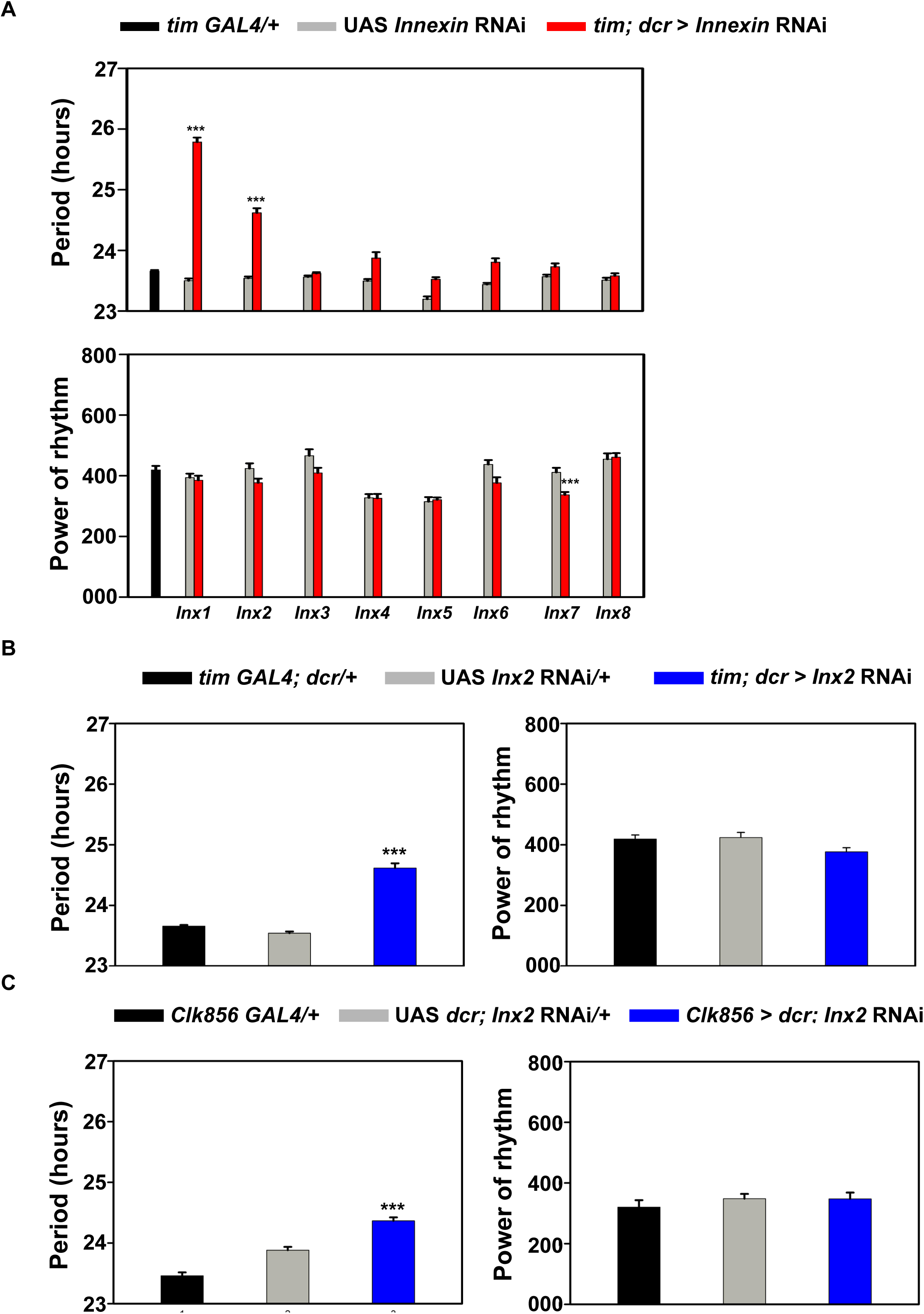
RNA interference screen of *Innexin*s in the clock neurons under DD 25 °C. **(A)** Mean free-running period (top) of flies with individual *Innexin* (*Innexin 1-8*) genes knocked down are being plotted along with their common *GAL4* control (*tim GAL4*/+) and respective UAS controls (UAS *Innexin* RNAi). Power of rhythm in case of individual knockdown of all the eight classes of *Innexins* along with their relevant parental controls are being plotted (bottom) (*n* > 26 flies for each genotype). (**B)** Mean free-running period (left) of experimental flies with *Inx2* knockdown using *tim GAL4* driver (*tim; dcr > Inx2* RNAi) is significantly longer than both driver control (*GAL4* control) and UAS control (left) while the power of the Chi-Square periodogram (right) of the experimental flies is not significantly different from the parental controls. 4 independent experiments were performed with similar results, with *n* > 22 flies for each genotype in each experiment. The graphs represent the results obtained from one representative experiment. **(C)** Mean free-running period (left) of experimental flies with *Inx2* knockdown using *Clk856* driver that targets most of the clock neurons (*Clk856* > *dcr*; *Inx2* RNAi) is significantly longer than parental controls and the power of rhythm (right) is not different from controls. 3 independent experiments were performed with similar results, with *n* > 25 flies for each genotype in each experiment. The graphs represent the results obtained from one representative experiment. Asterisks indicate significant difference between the experimental genotype and controls obtained using one-way ANOVA followed by Tukey’s HSD at *p<0.001*,error bars are SEM, period and power values are determined using Chi-square periodogram for a period of 7 days.

Additionally, we also knocked down *Innexin2* expression in all clock neurons using another driver, *Clk856 GAL4* which has a narrower expression pattern as compared to *tim GAL4*, but nevertheless targets most clock neurons except few DN3 (Gummadova et al., 2009). *Innexin2* knockdown using *Clk856 GAL4* also resulted in lengthening of free-running period by about an hour as compared to their respective parental controls (Fig. 1C, left). No significant difference in the power of the rhythm was observed in case of *Innexin2* knockdown as observed with the *tim GAL4* driver (Fig. 1C, right). We have also down-regulated the expression of *Innexin2* in the clock neurons using an alternate construct (BL 80409) on a different chromosome to account for the non-specific positional effects because of insertion of transgene. We observed similar extent of period lengthening with a different construct suggesting that the period lengthening phenotype seen is an effect of *Innexin2* knockdown and not because of positional effects (Supplementary fig. S1).

### *Innexin2* in ventral lateral neurons is important in determining the period of free-running rhythms

To determine whether *Innexin2* levels in specific subsets of the clock network modulates the free-running period, we used different drivers that target distinct subsets of circadian neurons; the ventral lateral neurons (*pdf GAL4*), ventral and dorsal lateral neurons (*dvpdf GAL4*), dorsal neurons (*Clk4.1M* and *Clk 4.5F GAL4*) as well as the glial cells (*repo GAL4*), the results of which are represented in the form of a table (Table 3). We found that *Innexin2* knockdown in lateral neurons with *dvpdf GAL4* and specifically the ventral lateral neurons using *pdf GAL4* resulted in lengthening of free-running period of the experimental flies as compared to controls (Fig. 2A, left), thus suggesting that *Innexin2* in these neurons are important in determining the period of the rhythm. As seen with the previous experiment, there was no difference in the power of the rhythm between the experimental and control flies (Fig. 2A, right). It is important to note here though that the period lengthening seen in case of experimental genotype when *Innexin*2 was knocked down with *pdf GAL4* (about 20 min) was not as long as that obtained using *tim GAL4* (about 60 min). This could be because of differences in the strength of *GAL4* drivers used or due to *Innexin*2 being functional in greater number of cells than the ones targeted by *pdf GAL4*. To examine the functional contribution of *Innexin2* in the ventral lateral neurons, we used *pdf GAL80* along with *tim GAL4* such that *Innexin2* expression is now down-regulated in all circadian neurons except the ventral lateral neurons. The efficiency of the *pdf GAL80* construct in suppressing *GAL4* expression in ventral lateral neurons was verified via immunohistochemistry using a GFP marker (Supplementary fig. S2).

**Table 3:** Effect of knockdown of *Innexin2* in smaller subsets of circadian pacemakers on free running rhythm properties. Table representing the average period (±SEM), power of the periodogram (± SEM) and % rhythmicity values of flies in case of *Innexin2* knockdown in different subsets of clock neurons and glia.

| Genotype | Period | Power of rhythm |
| --- | --- | --- |
| *pdf; dcr > Inx2 RNAi* | 23.65 ± 0.04 | 243.8 ± 10.4 |
| *Dvpdf > dcr; Inx2 RNAi* | 25.7 ± 0.16 | 234.4 ± 11.1 |
| *Clk4.1M > dcr; Inx2 RNAi* | 23.76 ± 0.04 | 358.8 ± 9.9 |
| *Clk4.5F > dcr; Inx2 RNAi* | 23.87 ± 0.04 | 372.9 ± 19.7 |
| *repo > Inx2 RNAi* | 23.9 ± 0.14 | 169.4 ± 12.8 |
| *w; pdfGAL4; +* | 23.26 ± 0.08 | 239.16 ± 7.89 |
| *yw; dvpdfGAL4; +* | 24.14 ± 0.04 | 295 ± 17.2 |
| *w; +; Clk4.1MGAL4* | 23.5 ± 0.05 | 359.2 ± 15.7 |
| *w; +; Clk4.5F GAL4* | 23.72 ± 0.06 | 294.6 ± 13.3 |
| *w; +; repo GAL4* | 23.7 ± 0.07 | 188.5 ± 11.8 |
| *yscv; +; Inx2 RNAi* | 23.65 ± 0.07 | 143.1 ± 5.57 |
| *w; dcr; Inx2 RNAi* | 23.8 ± 0.04 | 385.9 ± 18.29 |

**Figure 2:**
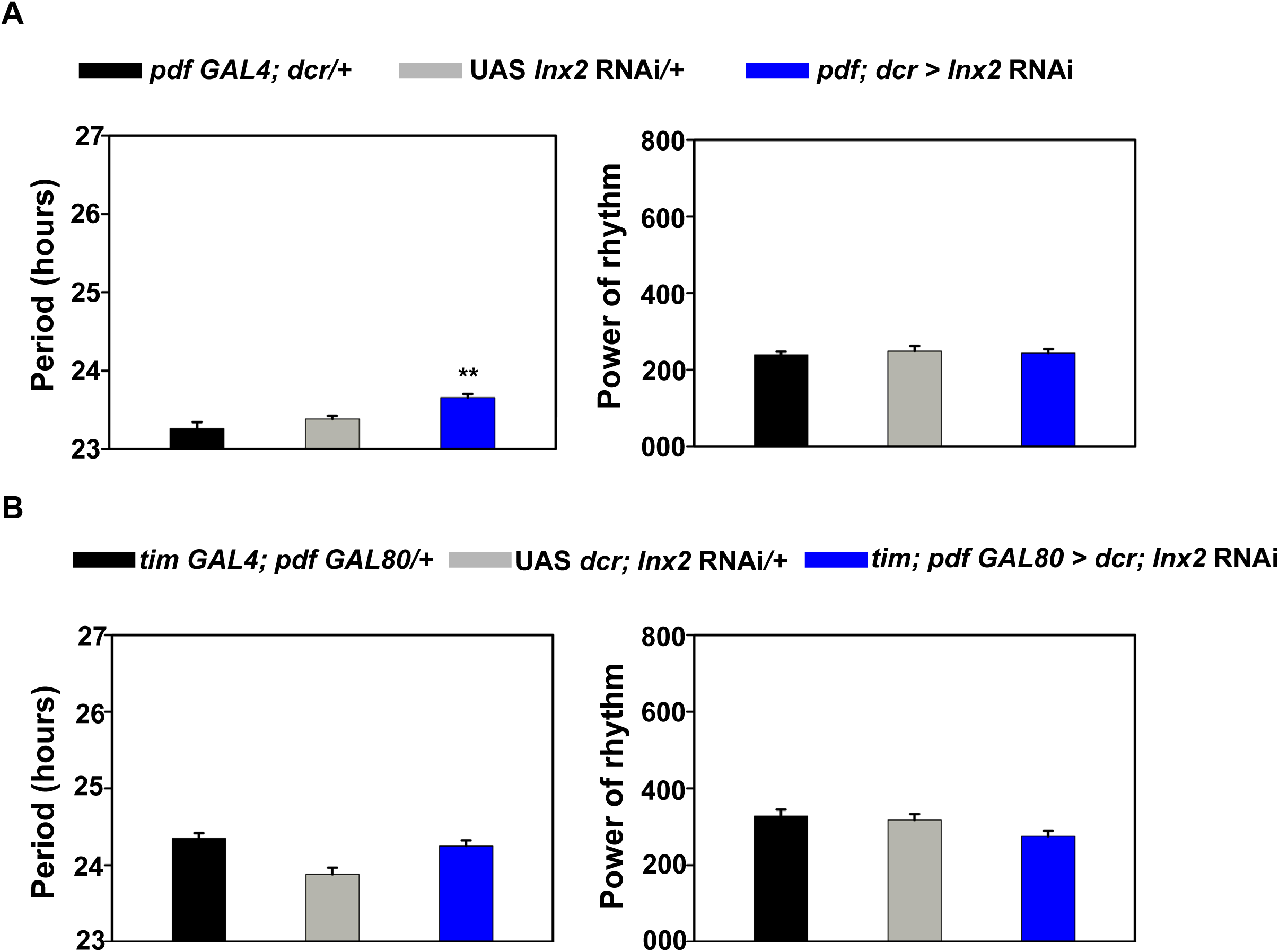
*Innexin2* knockdown in ventral lateral neurons lengthens free-running period. **(A)** Mean free-running period (left) of flies with *Innexin2* downregulated in the ventral lateral neurons (*pdf*; *dcr* > *Inx2* RNAi) is significantly longer than both its parental controls while the power of rhythm (right) is not different. 4 independent experiments were performed with similar results, with *n* > 28 flies for each genotype in each experiment. **(B)** Mean free-running period (left) of flies with *Innexin2* knockdown in all neurons of the clock circuit except the ventral lateral neurons (*tim*; *pdf GAL80* > *dcr*; *Inx2* RNAi) do not show a significant period lengthening as compared to its respective controls and the power of the rhythm (right) is not different between experimental flies and controls. 3 independent experiments were performed with similar results, with *n* > 25 flies for each genotype in each experiment. The graphs represent the results obtained from one representative experiment. Asterisks indicate significant difference between the experimental and control flies obtained using one-way ANOVA followed by Tukey’s HSD at *p < 0.01* (for fig. 2A), error bars are SEM, period values are determined using Chi-square periodogram for a period of 7 days.

*tim; pdf GAL80 > Inx2 RNAi flies* do not show a significantly lengthened period as compared to controls (Fig. 2B, left) and there was no change in power of the rhythm (Fig. 2B, right), suggesting that *Innexin2 function* in the ventral lateral subset contributes to free running period of activity rhythm.

### Distribution of INNEXIN2 in the circadian pacemaker circuit

Previous studies which have focused on the roles of *Innexin2* during nervous system development have found that during the larval stages, *Innexin2* is expressed to a large extent in the neural precursor cells and glial cells (Holcroft et al., 2013). To investigate the distribution of INNEXIN2 protein in the adult *Drosophila* brain and clock neuronal circuit, we performed immunohistochemistry using anti-INNEXIN2 antibody (Bohrmann & Zimmermann, 2008, kind gift from Prof. Michael Hoch, University of Bonn). To examine the co-localization of INNEXIN2 with the clock neurons, we used flies expressing nuclear localized GFP (GFP-NLS) with *tim GAL4* and co-stained the brain samples with antibody against INX2. In accordance with our behavioural results, we find that among clock neurons, INNEXIN2 is expressed only in the ventral lateral neuronal subset i.e. the small and large ventral neurons (Fig. 3). Additionally, we also consistently observe INNEXIN2 expression in about 8-9 cells in the dorsal side of the brain which are in close proximity to the dorsal DN1 neurons but do not co-localize with any of the dorsal clock neurons (Fig. 3, right, top and bottom). Thus, results from our behavioural and immunohistochemistry experiments put together suggest that INNEXIN2 is present and has functional roles in the ventral lateral neuronal subsets in determining the period of free-running rhythms.

**Figure 3:**
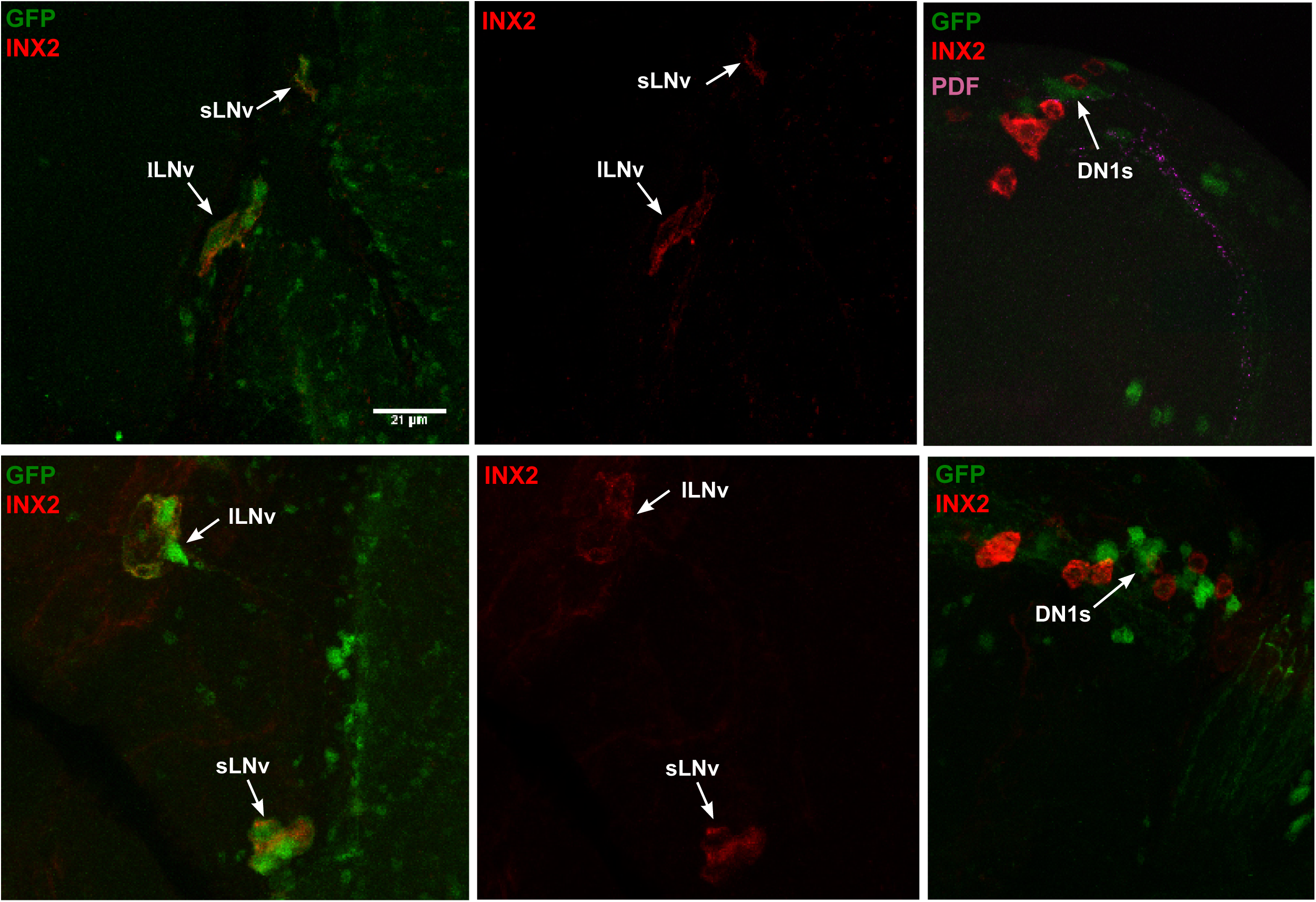
INNEXIN2 is localized to the small and large ventral lateral neurons in the circadian pacemaker circuit. Representative images from two different brains showing the distribution of INX2 protein among the clock cells. *tim* > *GFP-NLS* flies were stained with anti-GFP and anti-INX2 and checked for co-localization. INNEXIN2 was found to be predominantly localized to the membranes of small and large ventral lateral neurons (s-LNv and l-LNv) among the clock neurons (top and bottom, left and middle panels). INNEXIN2 was also present in 8-9 cells in the dorsal side of the brain in close proximity to DN1 neurons (top and bottom right panels). Brightness and contrast of representative images were adjusted in Fiji to facilitate better visualization. Arrows are used to indicate s-LNvs and l-LNvs, scale-bar represents 21 µm, *n* = 9 brain samples.

### Lengthening of free-running period due to *Innexin2* knockdown suggests its roles in mature adult circadian circuit

Since several previous studies have shown that *Innexin2* plays crucial roles during development of the fly of the nervous system, (Bauer et al., 2002, Bauer et al., 2004, Holcroft et al., 2013) we asked if the period lengthening seen in our experiments is due to defects in the development of the circadian pacemaker neuronal circuit or due to roles played by *Innexin2* in the mature, adult circuit. To distinguish between the two mechanisms, we temporally restricted the knockdown of *Innexin2* to the adult stages using the TARGET system (McGuire et al., 2004). All the flies used in this experiment were reared at a permissive temperature of 19 °C from embryonic stages till 3 days after eclosion to allow for the repression of *GAL4* by *tub GAL80*^*ts*^ and facilitate proper development including the final pruning of synaptic connections in the nervous system. The flies were then transferred to LD 12:12 at 29 °C and then assayed under constant darkness and restrictive temperature of 29 °C. The efficiency of the system in repressing the *GAL4* during development was verified by expressing UAS-*eGFP* under the same driver, *tim GAL4*; *tub GAL80*^*ts*^ and assessing GFP expression by immunohistochemistry in the larval stage L3 (Supplementary fig. S3). Significant lengthening of period of activity rhythm was observed in experimental flies as compared to controls even when *Innexin2* knockdown in clock neurons was restricted to adult stages (Fig. 4A, left), suggesting that *Innexin2* plays a role in the adult circadian circuit to determine the period of free-running rhythms. However, we acknowledge the fact that in the larval stages, we see faint GFP staining in 1 out of 3-4 s-LNvs/hemisphere in some of the brain samples, thus suggesting that the *tub GAL80*^*ts*^ was unable to completely repress the expression of *Innexin2* RNAi during development. But to a large extent, we can conclude that the period lengthening that we observe are largely due to its roles in the adult circuit although *Innexin2* could possibly have some roles in the development of the circuit. In this specific case, the power of rhythm of the experimental flies was found to be significantly higher than the controls (Fig. 4A, right). We also restricted the knockdown to the ventral lateral neurons in the adult stages only using *pdf GAL4* and *tub GAL80*^*ts*^. In this case, however, we observed a significant lengthening of period from only one parental control (Fig. 4B, left), although we see a trend towards longer period values. This could be because *Innexin2* levels modify development of these cells such that the period lengthening seen during constitutive knockdown is an additive effect of its roles during development and in the mature circuit. In this case too, we found that the power of the rhythm is significantly higher than controls (Fig. 4B, right).

**Figure 4:**
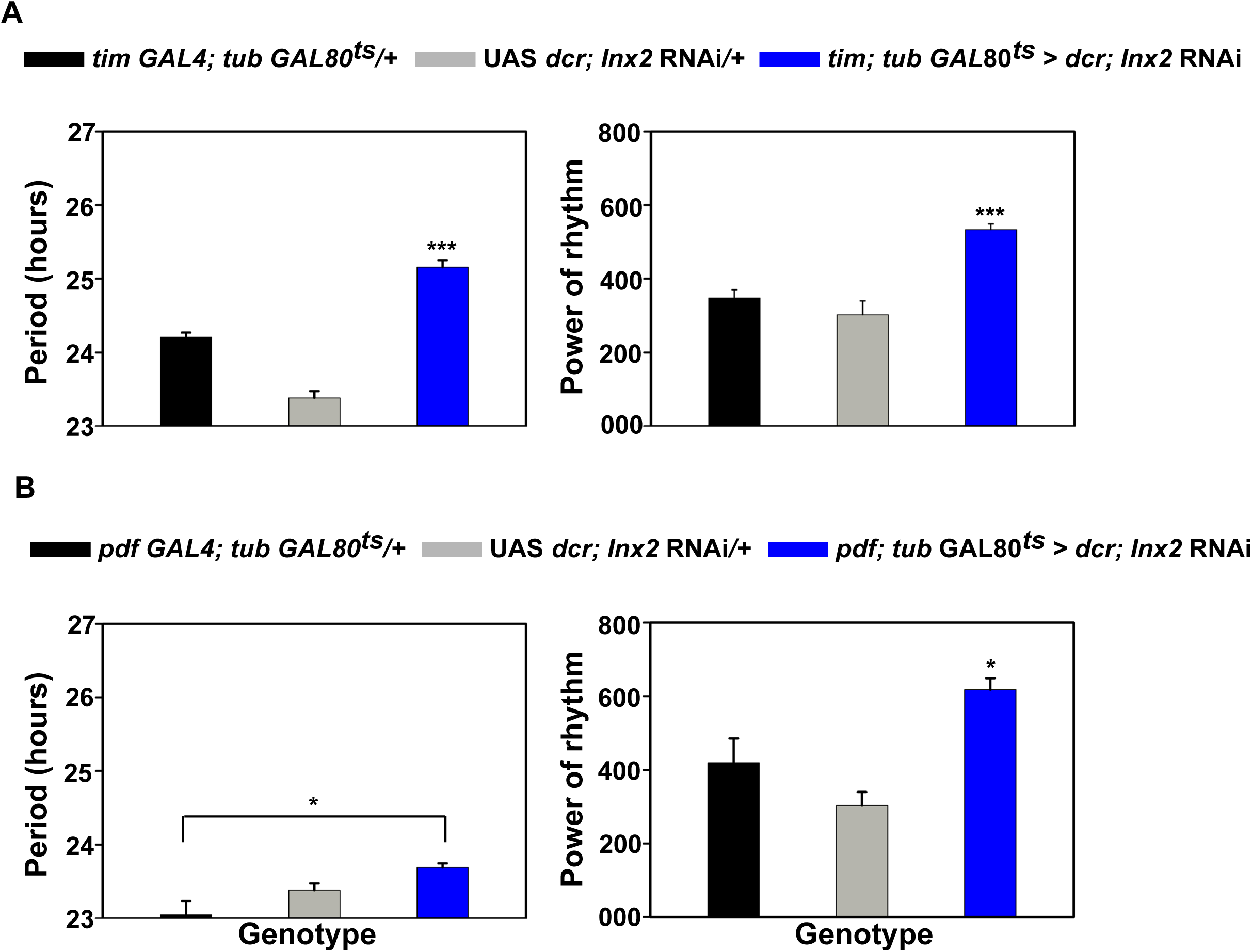
Period lengthening seen in case of *Innexin2* knockdown is not due to developmental defects. **(A)** Mean free-running period (left) of activity rhythm when *Innexin2* knockdown in all the clock neurons is restricted to adult stages (*tim*; *tub GAL*80^ts^ > *dcr*; *Inx2* RNAi), is significantly longer compared to its relevant parental controls. Power of rhythm (right) of the experimental flies are found to be significantly higher than controls (*n >* 21). **(B)** Mean free-running period (left) of activity rhythm in case of adult-specific knockdown of *Innexin2* only in the ventral lateral neurons (*pdf*; *tub GAL*80^ts^ > *dcr*; *Inx2* RNAi) was only significantly different from its *GAL4* control. Power of the rhythm (right) was not different from controls (*n* > 15). Asterisks indicate significant difference between the experimental flies and controls obtained using one-way ANOVA followed by Tukey’s HSD at *p<0.001* (for panel A), *and p<0.05* (for panel B), error bars are SEM, period and power values are determined using Chi-square periodogram for a period of 10 days in DD 29° C.

### Knockdown of *Innexin2* delays the phase of PERIOD protein oscillation in the circadian clock network and leads to higher levels of PDF accumulation in s-LNv dorsal terminals

In order to examine the mechanism by which *Innexin2* influences the period of free-running rhythms in *Drosophila*, we tracked the oscillation of the core molecular clock protein, PERIOD (PER) in the 6 circadian pacemaker cell clusters and also the levels of the clock output neuropeptide PDF in the dorsal projections over a 24-h cycle on day 3 of constant darkness (DD) in both control (*dcr*; *Inx*2 RNAi; UAS control) and experimental (*Clk856* > *<u>dcr</u>*; *Inx*2 RNAi) flies. We found that even though *Innexin*2 is present only in the ventral lateral neurons, the oscillation of PERIOD protein is phase-delayed in most clock neurons in the circuit. Using a COSINOR-based curve-fitting method, we found a significant 24-h rhythm in PER oscillation in s-LNv in case of both control and experimental flies (Fig. 5, Table 4). The phase of PER oscillation in case of experimental flies was however significantly delayed from control flies (Fig. 6A) suggesting that *Innexin2* knockdown results in a shift in the core molecular clock oscillation. The amplitude of oscillation was not found to be different from controls (Fig. 6B). In case of PER oscillation in l-LNv, we could detect a significant 24-h rhythm in case of both control and experimental flies (Fig. 5, Table 4). The phase of the oscillation was also significantly delayed in case of experimental flies as compared to controls (Fig. 6A). Even though the amplitude of oscillation in the l-LNv in experimental flies was not found to be different from the controls (Fig. 6B), the amplitude of oscillation in l-LNv in control flies was found to be significantly lower than that of s-LNv (Supplementary fig. S4). In case of 5^th^ s-LNv, we observe a significant 24-h oscillation in the control flies. But in case of experimental flies, no significant rhythmicity was detected using the COSINOR-based method (Fig. 5, Table 4). In case of LNd, both the control and experimental flies show a significant 24-h rhythm in PER oscillation (Fig. 5, Table 4). The phase of oscillation of experimental flies was found to be significantly delayed from the controls (Fig. 6A). The amplitude of oscillation in LNd was not found to be different between the control and experimental flies (Fig. 6B), whereas the amplitude of oscillation of control flies was found to be significantly lower as compared to s-LNv (Supplementary fig. S4)

In case of DN1, although we could detect a significant 24-h periodicity in control flies, the amplitude of the oscillation was highly dampened. (Fig. 5, Table 4) In experimental flies, however, DN1s do not show significant rhythmicity in PER oscillation and had overall low amplitude (Fig. 5, Table 4). In case of DN2s, both control and experimental flies do not show a significant 24-h rhythmicity and have highly dampened oscillation (Fig. 5, Table 4). Since PDF is an important neuropeptide in the circadian pacemaker circuit which plays a role in the synchronization of all the neurons and PDF levels in dorsal projections cycle with a 24-h periodicity under DD, we examined whether *Innexin*2 knockdown has an effect on the PDF levels or oscillations in the s-LNv dorsal terminal. We found that, both control and experimental flies show a robust 24-h oscillation in PDF levels in the dorsal projections (Fig. 7A). In contrast to PER oscillation, there was no significant difference in the phase of PDF oscillation in experimental flies as compared to the control (Fig. 7B, left, Table 4). However, we observed that PDF levels and amplitude was significantly higher in experimental flies as compared to the control (Fig. 7B, right, Table 4).

**Table 4:** Rhythm properties of PERIOD expression in distinct cell groups of control and experimental genotypes. Table representing the parameters obtained after fitting a COSINE curve on the PER and PDF intensity data obtained over a 24-h period for all the circadian neuronal subsets on third day of constant darkness. The parameters obtained are p-values and percent rhythm (PR) to test for significant 24-h periodicity, and the amplitude and phase values along with their respective standard errors (SE).

| Cell type | Period | p value | PR | Amplitude ± SE | Phase ± SE |
| --- | --- | --- | --- | --- | --- |
| s-LNv control | 24-h | 1.38e-14 | 62.53 | 32.62 ± 3.13 | -320.67 ± 5.76 |
| s-LNv experimental | 24-h | 4.12e-13 | 61.34 | 24.81 ± 2.54 | -45.89 ± 6.66 |
| l-LNv control | 24-h | 7.57e-05 | 24.98 | 10.68 ± 2.28 | -297.50 ± 13.52 |
| l-LNv experimental | 24-h | 6.85e-05 | 27.35 | 11.51 ± 2.44 | -75.76 ± 12.83 |
| 5<sup>th</sup> s-LNv control | 24-h | 0.005 | 26.78 | 28.18 ± 8.49 | -311.54 ± 13.3 |
| 5<sup>th</sup> s-LNv experimental | 24-h | 0.36 | 5.09 | 7.70 ± 5.79 | -0.06 ± 33.28 |
| LNd control | 24-h | 9.89e-05 | 30.84 | 19.11 ± 4.06 | -306.95 ± 10.85 |
| LNd experimental | 24-h | 4.54e-05 | 29.16 | 14.19 ± 2.9 | -43.72 ± 13.89 |
| DN1 control | 24-h | 0.007 | 17.86 | 9.88 ± 3.01 | -257.73 ± 15.68 |
| DN1 experimental | 24-h | 0.15 | 6.42 | 6.51 ± 3.32 | -52.54 ± 33.44 |
| DN2 control | 24-h | 0.97 | 0.16 | 0.86 ± 3.89 | -44.09 ± 214.23 |
| DN2 experimental | 24-h | 0.73 | 1.61 | 2.21 ± 2.9 | -25.65 ± 86.24 |
| PDF in DP control | 24-h | 1.38e-07 | 34.01 | 1.16 ± 0.18 | -51.30 ± 8.7 |
| PDF in DP experimental | 24-h | 6.25e-13 | 58.44 | 2.30 ± 0.24 | -64.10 ± 6.3 |

**Figure 5:**
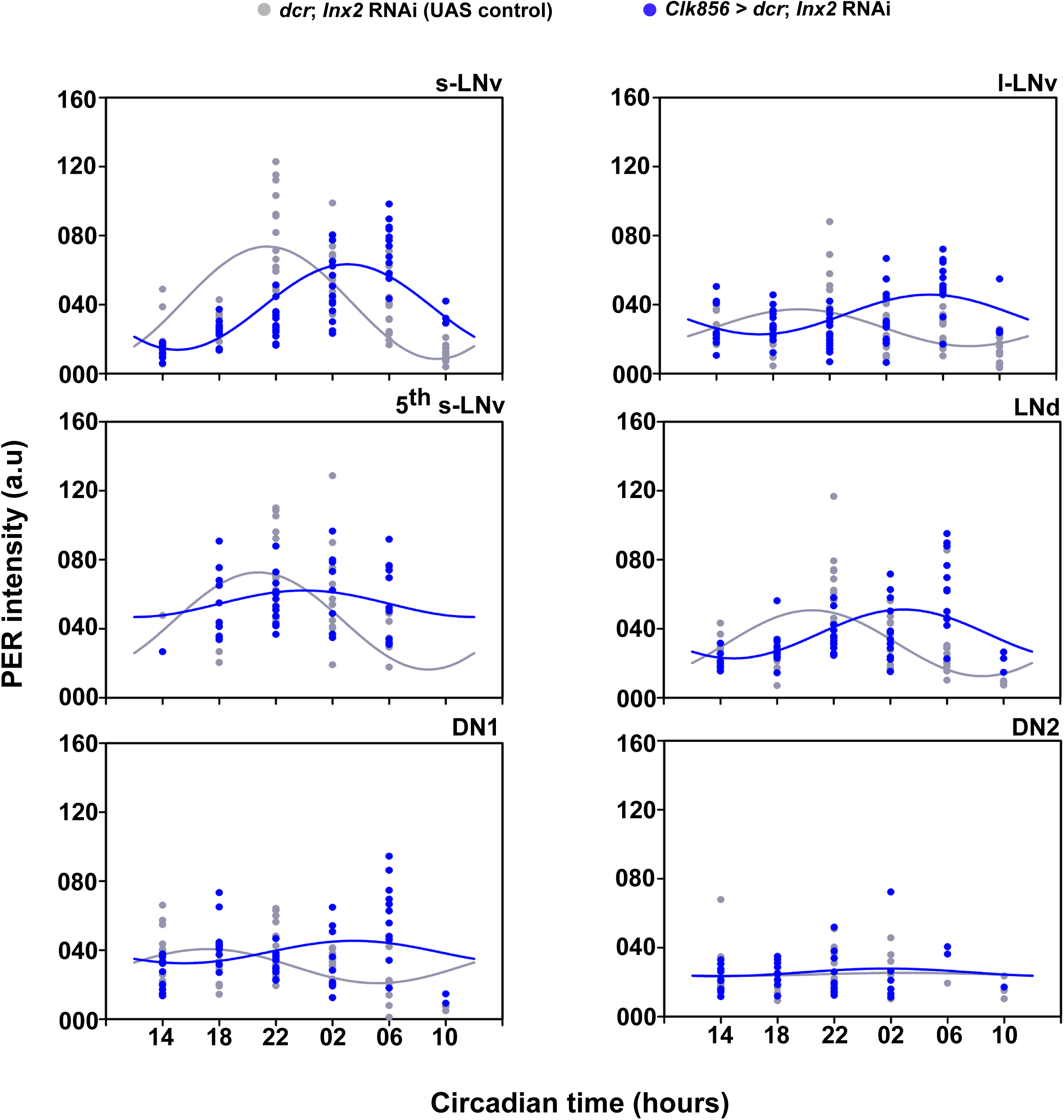
Knockdown of *Innexin2* delays the oscillation of PER in most clock neuronal subsets in the circadian pacemaker circuit. Scatter plots of PER staining intensities in each of bilaterally located six distinct neuronal clusters of the circadian pacemaker network of both the control (*dcr*; *Inx2* RNAi) and experimental (*Clk856* > *dcr*; *Inx2* RNAi) flies plotted at different time-points over a 24-h cycle on third day of DD. Each dot represents the mean PER intensity value averaged over both the hemispheres of one brain. The gray and blue lines are the best fit COSINE curve from the parameters that were extracted from the COSINOR analysis. *n* > 11 brain samples both in case of control and experimental flies for all time points except for CT10 in case of experimental flies where a technical difficulty resulted in n =3 brain samples only being imaged. In case of 5^th^ s-LNv, we could not detect any cells at CT10 in experimental samples, hence that time point was not included for analysis.

**Figure 6:**
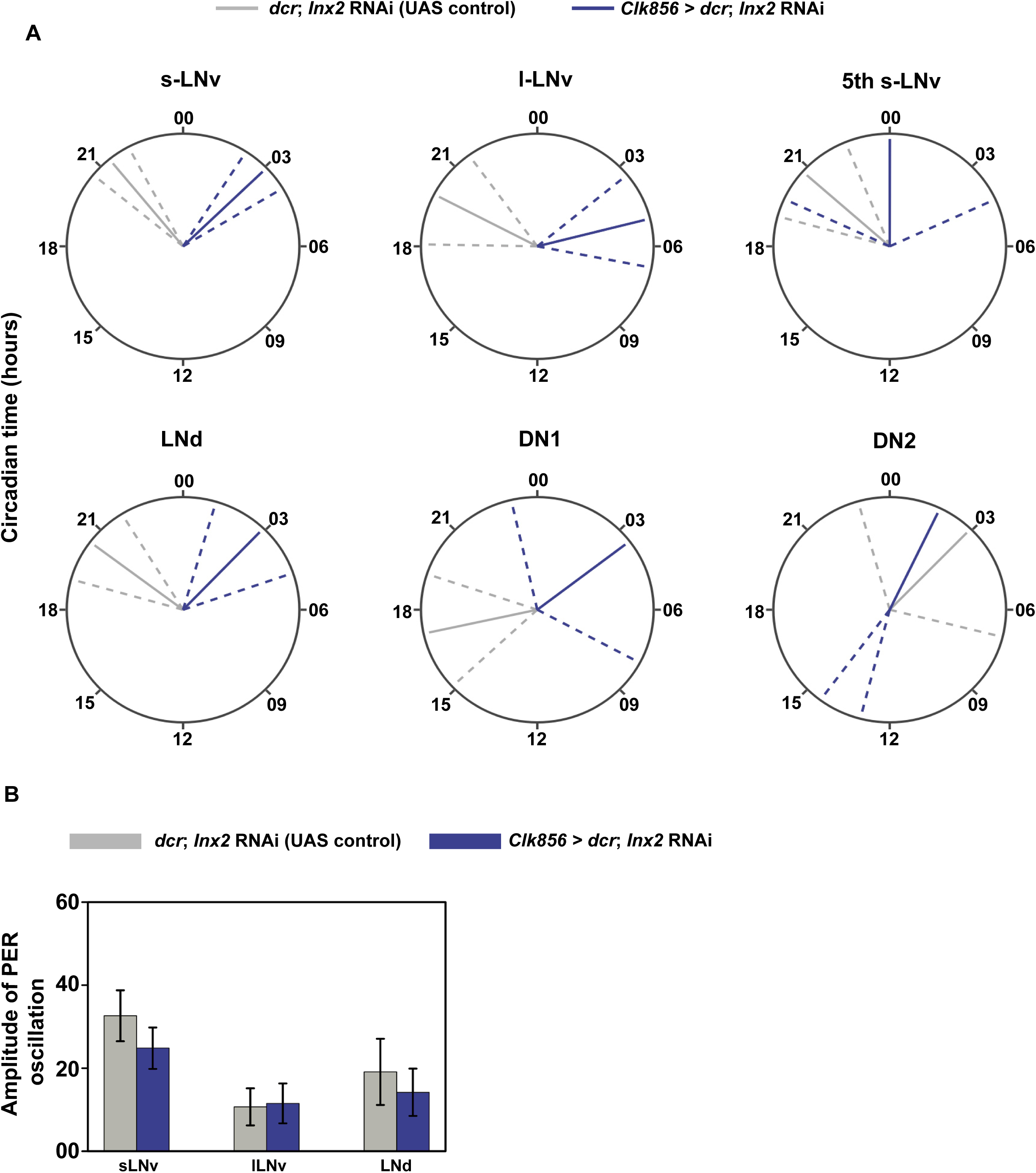
*Innexin2* knockdown affects the phase but not the amplitude of PER oscillations in the clock cells. **(A)** Polar-plots depicting the acrophase of PER oscillation in control (*dcr*; *Inx2* RNAi) (gray lines) and experimental flies (*Clk856* > *dcr*; *Inx2* RNAi) (blue lines) in all six distinct neuronal clusters of the circadian pacemaker network. The acrophase values obtained after COSINOR curve fitting are shown as solid lines and the error (95% CI values) is depicted as dashed lines around the mean for all the cell types. Non-overlapping error values indicate that phase values of experimental flies are significantly different from controls as seen in the case of s-LNv, l-LNv, and LNd. Although the experimental lines in case of DN1 have non-overlapping phase value from controls, no significant 24-h rhythms were detected in this case. **(B)** Amplitude values obtained from COSINOR curve fits are plotted for control and experimental flies for those cell groups which show significant 24-h rhythms (s-LNv, l-LNv and LNd) and the error bars represent 95% CI values. Overlapping error bars indicate that amplitude values of experimental flies are not significantly different from controls. See Table 4 for more details.

**Figure 7:**
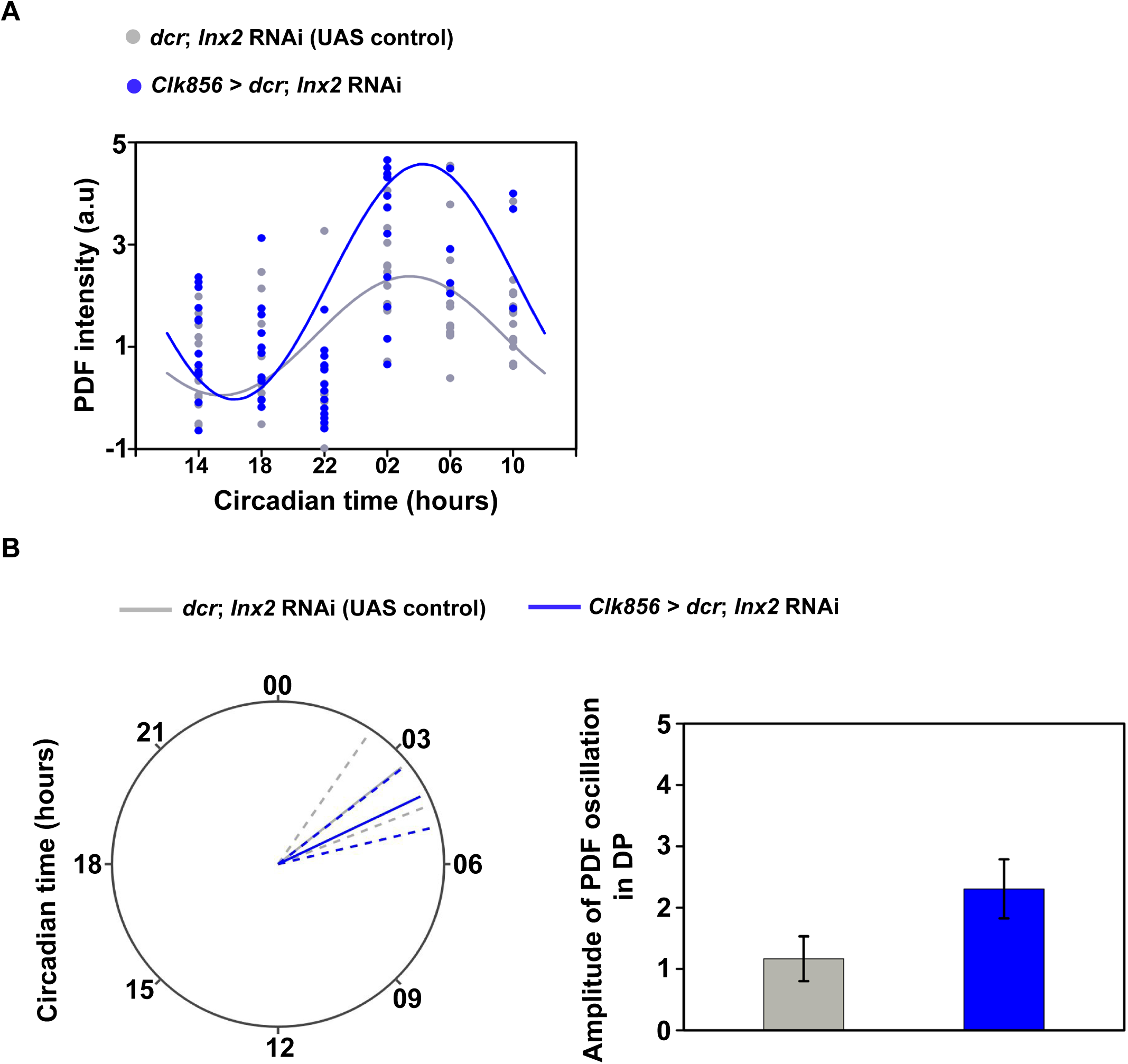
*Innexin2* knockdown affects the amplitude of PDF oscillations in the sLNv dorsal projections. **(A)** Scatter plots of PDF intensity in the s-LNv dorsal projection of both the control (*dcr*; *Inx2* RNAi) and experimental (*Clk856* > *dcr*; *Inx2* RNAi) flies plotted at different time-points over a 24-h cycle on third day of DD. Each dot represents the mean PDF intensity value averaged over both the hemispheres of one brain. *n* > 10 brain samples both in case of control and experimental flies for all timepoints except for CT10 in case of experimental lines where a technical difficulty resulted in n =3 brain samples only being imaged. The gray and blue lines are the best-fit COSINE curve from the parameters that were extracted from the COSINOR analysis. **(B)** Polar-plots depicting the acrophase of PDF oscillation in control (*dcr*; *Inx2* RNAi) and experimental flies (*Clk856* > *dcr*; Inx2 RNAi) (left). The mean phase values obtained after COSINOR curve fitting are shown as solid lines and the error values (95% CI) are depicted as dashed lines around the mean. Overlapping error bands indicate that the experimental phases are not significantly different from controls. Amplitude values obtained from COSINOR curve fits are plotted for control and experimental flies and the error bars represent 95% CI values (right). Non-overlapping error bars indicate that experimental lines are significantly different from controls. See Table 4 for more details.

## Discussion

The neuronal and molecular mechanisms underlying circadian rhythms have been extensively studied in *Drosophila melanogaster* for many years now. Yet, an interesting question that remains to be answered is how are free-running behavioural rhythms with a near 24-h periodicity generated by the network. Although membrane excitability states have been shown to be important for circadian behaviour in *Drosophila* and there are some reports from mammalian studies on the role played by gap junctions in SCN neural network, there have been no systematic studies investigating the importance of electrical synapses in the circadian pacemaker circuit in *Drosophila*. We report here for the first time that gap junction proteins play important roles in the *Drosophila* circadian pacemaker circuit to influence the period of free-running rhythms.

Our screen revealed that gap junction genes *Innexin1* and *Innexin2* may play important roles in determining the period of free-running rhythms. We carried out further experiments to understand the mechanism by which *Innexin2* modulates the free-running period. We found that among the clock neurons *Innexin2* is present and functions solely in the small and large ventral neuronal subsets. Several previous studies have shown the importance of s-LNv and PDF under constant darkness to generate free-running rhythms of near 24-h periodicity (Renn et al., 1999, Grima et al., 2004, Stoleru et al., 2004, Park et al., 2000, Yoshii et al., 2009), although, some studies challenge the notion of the hierarchical role played by s-LNv in the network and instead show it to be composed of multiple coupled oscillators with each of them contributing to the resultant free-running period in behaviour (Sheeba et al., 2008, reviewed in Sheeba, Kaneko, et al., 2008, Yao & Shafer, 2014, Dissel et al., 2014, Schlichting et al., 2019, Delventhal et al., 2019). In either case, it is well-accepted now that s-LNv play significant role in determining the period of the network. Since, *Innexin2* is present in the s-LNv and influences the free-running period, we investigated the underlying mechanism. As a first step towards the same, we examined the oscillation of molecular clock protein PERIOD on third day in DD in both control and experimental flies in which *Innexin2* was knocked down in all clock neurons. To our surprise, we found that the molecular clock is delayed in most clock neurons even though *Innexin2* was only found to be present in the LNvs. Phase of PER oscillations in case of s-LNv, l-LNv and LNd cell types in experimental flies were found to be significantly delayed as compared to control flies. Although, amplitude of oscillations were not different between the experimental and control flies in each of these cell types, amplitude of oscillations in l-LNv and LNd in control flies were significantly lower than the s-LNv. It has been observed in several previous studies that amplitude of PER oscillation in l-LNv dampen after 2 days in constant darkness (Yang & Sehgal, 2001, Shafer et al., 2002, Peng et al., 2003, Roberts et al., 2015) and we observe something similar in our experiment. In case of LNd, a previous study has shown them to be a heterogenous group of cells which are differentially coupled to sLNv with some having stronger and others having weaker coupling (Yao & Shafer, 2014). However, in this experiment, we had no means to distinguish amongst them and have averaged PER intensities across all the 5-6 cells which could also contribute to the observed low amplitude values. 5^th^ s-LNv shows rhythmic PER oscillations in control flies whereas the experimental flies show arrhythmicity. This lack of rhythmicity in 5^th^ s-LNvs in experimental flies could be explained in the context of a previous study which shows that 5^th^ s-LNv being only weakly coupled to PDF^+^ s-LNv and also receives input from PDFR^−^ LNd (Yao & Shafer, 2014), and the arrhythmicity observed could be because of conflicting signals received from the two cell clusters of different periodicities. Similarly, in case of DN1, we observe highly dampened rhythms in control flies, similar to previous studies which have reported less robust rhythms, with patterns of dampening amplitude and loss of coherent rhythmicity in DN1 over 6 days in DD (Roberts et al., 2015, Yoshii et al., 2009). Lack of rhythmicity in DN1 in case of experimental flies could be could be explained by the fact that these cells receive conflicting signals from sLNv and LNd with different periodicities (Zhang et al., 2010). In case of DN2, both the control and experimental flies do not show a significant 24-h rhythmicity and have highly dampened rhythms. Although PDFR is expressed in DN2, molecular clock in DN2 was found to be independent of the control of sLNv and do not seem to have profound effects on rhythmic activity-rest behaviour in DD (Stoleru et al., 2005).

How does a gap junction protein localized to the membrane of LNv affect the phase of molecular clock protein oscillations in the network? It is possible that INNEXIN2 present in the membrane of LNv affects the membrane excitability state of these neurons which then affects the core molecular clock. Several previous studies have shown that membrane excitability states of the LNv can affect both the core molecular clock and features of activity-rest rhythms. Both small and large LNvs show time-of-day dependence in membrane electrical activity such that it is depolarized in the early part of the day and becomes hyperpolarized in the later part of day (Sheeba, Gu, et al., 2008, Cao & Nitabach, 2008). Constitutive hyperexcitation of LNv membrane by expression of sodium channel NaChBac results in complex rhythms with multiple periodicities in behaviour along with desynchronization of molecular clocks in the circuit and disrupted cycling of PDF in dorsal terminals (Nitabach et al., 2006). Silencing of LNv by expressing the inward potassium rectifier channel Kir2.1 results in behavioural arrhythmicity, and disruption of molecular clocks (Nitabach et al., 2002), athough adult-specific silencing of these neurons have milder effects (Depetris-Chauvin et al., 2011). Membrane excitability states can also affect the transcriptional states of LNv such that hyperexcitation can generate a very different gene expression profile in the cell as compared to hyperpolarization (Mizrak et al., 2012). Thus, it is evident that membrane excitability states can affect the core molecular clock and the delay in PER oscillations observed in case of *Innexin2* knockdown could be via changes in membrane excitability states of the LNv. This is in contrast to what has been reported about the mechanism by which gap junction gene *Cx36* functions in the SCN which states that although *Cx36* affects the synchrony of firing of SCN cells and period and amplitude of behavioural rhythms, it does not affect the period, amplitude or synchrony of molecular clocks in SCN slices (Long et al., 2005, Wang et al., 2014, Diemer et al., 2017).

The mechanism of action of gap junctions have been well-studied in case of *Connexins* and not so much in case of *Innexin*s, although the basic structure and function of the two are comparable (Beyer & Berthoud, 2018). Gap junctions in nervous systems are known to facilitate generation and modulation of synchronous firing of neurons which can also affect the release of neuropeptide and neurotransmitters. Other than being involved in electrical coupling, gap junctions also facilitate passage of secondary messengers and small molecules whose size is less than 1 kDa between neuron-neuron or they can act as hemichannels and facilitate transport between neurons and extra cellular matrix (reviewed in Nielsen et al., 2012).

We propose two possible hypotheses for the mechanism by which *Innexin2* in LNv membrane could affect the molecular clock oscillations in circadian neurons. The first possibility is that *Innexin2* could be involved in electrical coupling and synchronous neuronal firing among the LNv and the absence of *Innexin2* results in desynchronized firing which could be reflected in the release of PDF. In our experiments, we observe that the amplitude of PDF oscillation and level of PDF in the dorsal projections is much higher in experimental flies than in control ones. Several studies have shown that PDF acts to lengthen the period of the circadian network and that overexpression or ectopic expression of PDF in the dorsal protocerebrum lengthens the period of activity rhythms, leads to desynchronization of activity-rest behaviour and molecular clocks in the circadian neurons (Charlotte Helfrich-Förster et al., 2000, Wülbeck et al., 2008, reviewed in Shafer & Yao, 2014). Thus, *Innexin2* may play a role in synchronized firing and release of PDF in the dorsal projections, the absence of which could result in PDF being present in the projections at a higher level and could delay the molecular clocks in all the other clock neurons. The alternate possibility could be that *Innexin2* is necessary to maintain a certain absolute value of membrane potential of the LNv at certain times of the day or are important to regulate the number, frequency or pattern of action potential firing in these cells and a disruption in this process in case of *Innexin2* knockdown translates into a delay in the molecular clocks in LNvs which is then transmitted to other neurons in the circuit via PDF. Indeed, there is some evidence to suggest that gap junctions affect the frequency of firing of action potentials in the l-LNv membrane. An experiment performed by Cao and Nitabach using gap junction blocker Carbenoxolone in the bath and recording action potentials from the l-LNv shows that blocking electrical synapses reduces the frequency of firing of action potentials in these cells (Cao & Nitabach, 2008). At this time we do not know how these changes in firing frequency may alter the core molecular clock in circadian pacemakers, however studies performed on neurons from the dorsal root ganglion suggest that expression levels of many genes are highly affected by firing frequency (Fields et al., 1997, Lee et al., 2017).

While current experiments do not allow us to distinguish between the above mentioned possibilities, further experiments need to be done to address the following questions. Do *Innexin2* mutant flies exhibit time-of-day dependent oscillations in membrane potential in the LNv? Does knockdown of *Innexin2* affect the absolute membrane potential value of LNv as compared to controls or does it affect the synchronous firing of those neurons? These questions can be addressed by electrophysiological recording of the LNv although recording from the s-LNv is technically challenging. An alternate approach would be to measure the spontaneous calcium activity rhythms in these cells and all the other cells of the circuit in a time-of-day dependent manner which can reveal any changes in membrane electrical state in the circadian pacemaker neurons (Liang et al., 2016).

Additionally, we also find that *Innexin2* plays predominant roles in the mature, adult circadian circuit to influence the period of free-running rhythms although our results also suggest that it could have some roles to play in the development of clock neuronal circuit. Also, in case of adult specific knockdown of *Innexin2* in all clock neurons as well as ventral lateral neurons, we observe that power of rhythm in experimental flies was much higher than that of controls which we do not observe in case of constitutive knockdown of *Innexin2* during both developmental and adult stages. One probable explanation could be that the circuits are wired differently in the presence and absence of *Innexin*s such that instead of multiple oscillators with different periods contributing to the behaviour, the relative contribution of some oscillators have increased or decreased. However, further experiments are needed to validate the same.

How does *Innexin2* form gap junctions in the LNv? Previous studies have reported that *Innexin2* can form functional heterotypic gap junctions with *Innexin1, Innexin3* or *Innexin4* as well as form homotypic gap junctions with itself to facilitate passage of ions and/or secondary messengers or small molecules (Bauer et al., 2003, Bohrmann & Zimmermann, 2008, Holcroft et al., 2013). Alternatively, *Innexin2* can also function as hemichannels and facilitate coupling between the cell and extracellular matrix, thus allowing passage of ions or small molecules. Since, we do not observe any period lengthening in case of *Innexin3* or *Innexin4* knockdown, the only possibilities are *Innexin2* forming functional heterotypic gap junction with *Innexin1* or homotypic gap junctions with itself. With results from this manuscript and our results from another study (Ramakrishnan and Sheeba, *manuscript in preparation*) put together, we can conclude that *Innexin2* either forms homotypic gap junctions or functions as hemichannels in s-LNv, and in case of l-LNv, *Innexin2* and *Innexin1* probably form heterotypic gap junctions. However, this needs further validation. Recently, a technique was developed by Wu et al to demonstrate functional electrical coupling among cells called PARIS (Pairing Actuators and Receivers to Optically Isolate Gap junctions) (L. Wu et al., 2019). Studies to determine functional electrical coupling among the LNvs using this technique both in control flies as well as *Innexin1* and *Innexin2* mutants to identify the type (homotypic or heterotypic) of channels present in these cells would be valuable Thus, our findings highlighting a hitherto unknown role for *Innexins* in the adult circadian pacemaker circuit of *D.melanogaster* in determining free-running period of activity rhythms via influencing a core clock protein PER within s-LNv and l-LNv reveals the action of a combination of electrical and chemical synapses in circadian pace-making.

## Figure legends

**Supplementary fig. S1: Knockdown of Innexin2 in all clock neurons using an alternate construct (BL 80409) lengthens the period of free-running rhythms** Mean free-running period (left) of flies with *Innexin2* downregulated in all clock neurons (*Clk856 > dcr; Inx2 RNAi*) is significantly longer than both its parental controls while the power of rhythm (right) is significantly lower than only one parental control. *n* > 15 flies for each genotype.

**Supplementary fig. S2: Verification of the efficiency of *pdf GAL80* construct** The efficiency of *tim*; *pdf GAL80* construct was verified by crossing *tim GAL4*; *pdf GAL80* flyline with *GFP-NLS*, dissecting the brains at ZT22 and staining with GFP and PER. Both small and large LNvs did not show any GFP staining (left), whereas strong PER staining was observed in all clock neurons (middle). n=6 brain samples were imaged.

**Supplementary fig. S3: Verification of the efficiency of *tub GAL80***^***ts***^ **construct** The efficiency of *tim; tub GAL80*^*ts*^ construct was verified by crossing *tim; tubGAL80*^*ts*^ with *eGFP*. The flies were reared at a permissive temperature of 19 °C. Larvae (L3 stage) from the progeny was dissected and stained with GFP and PER antibodies. s-LNv do not show presence of GFP in permissive temperatures (left), whereas strong PER staining was observed in this cell (middle). n=5 brain samples were imaged.

**Supplementary fig. S4: Amplitude of PER oscillations are different in circadian neuronal neuronal subsets on third day of DD** Amplitude values of PER oscillation obtained from COSINOR curve fits are plotted for control (UAS *dcr; Inx2 RNAi*) flies for all the circadian neuronal subsets. The error bars represent 95% CI values. Non-overlapping error bars in case of l-LNv, LNd and DN1 compared to s-LNv indicate that these amplitude values are different from s-LNv.

## Acknowledgements

We thank JNCASR Intramural funds and a grant from DST-SERB (CRG/2019/006802) for funding this work. We thank DST-INSPIRE for providing fellowship to AR. We thank Prof. Michael Hoch, Dr. Reinhard Bauer and Prof. Jeffrey Hall for generously sharing anti-INX2 (guinea pig) and anti-PER (rabbit) antibody respectively, and Orie Shafer, Todd Holmes, Michael Rosbash and Charlotte Helfrich-Forster for kindly sharing fly lines, Jaimin Bhatt for help with dissections for immunohistochemistry experiment, Mr. Prajwal and Ms. Suma for help with imaging and Rajanna and Muniraju for technical assistance. We are extremely grateful to Abhilash Lakshman for useful discussions regarding statistics for immunohistochemistry experiment data, for help with running the CATCosinor function from the CATkit package and writing custom codes in Plotly for plotting polar plots in R. We also thank Prof. Gaiti Hasan for useful discussions and inputs at various stages of this project.

