## Supplementary files for "Gap junction protein INNEXIN2 modulates the period of free-running rhythms in *Drosophila melanogaster*"

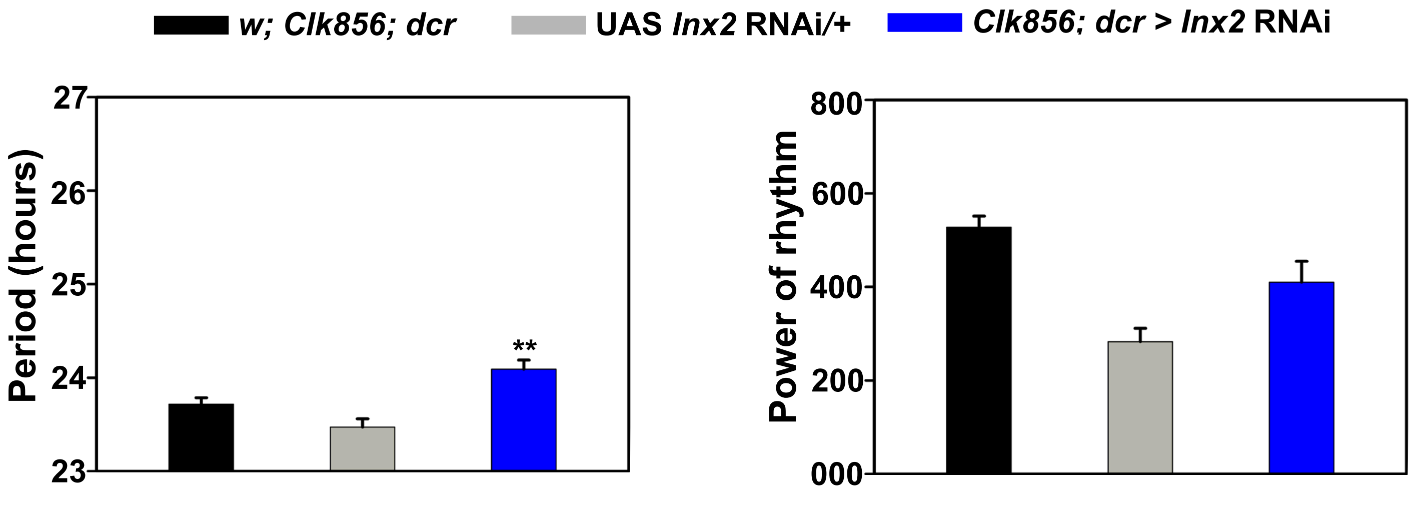


**Supplementary fig. S1:** Knockdown of *Innexin2* in all clock neurons using an alternate construct (BL 80409) lengthens the period of free-running rhythms.

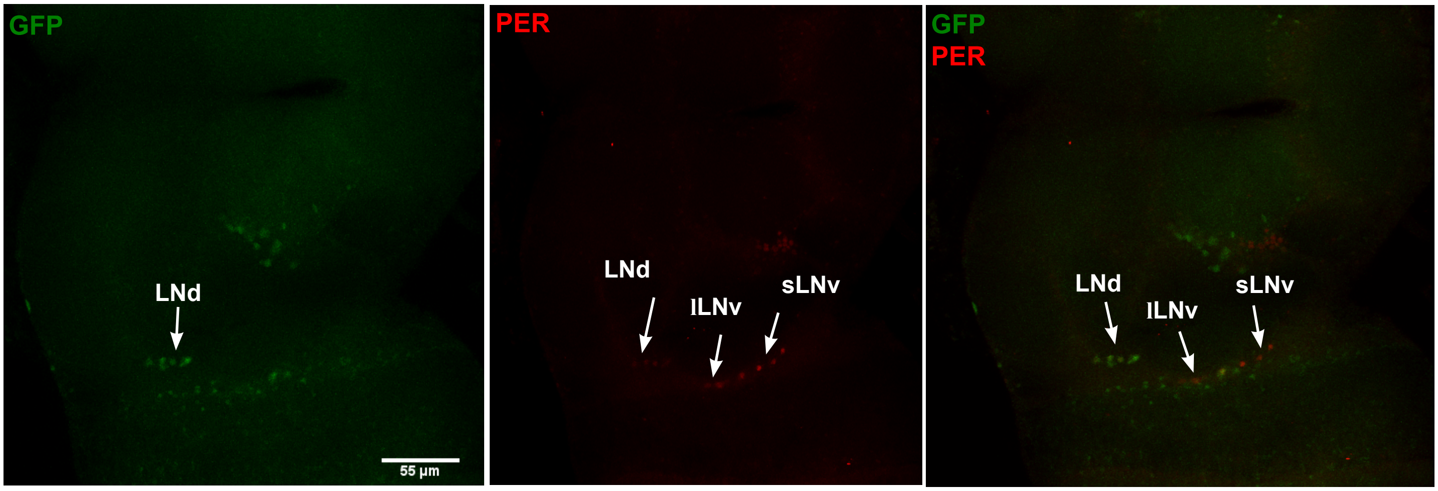


**Supplementary fig. S2:** Verification of the efficiency of *pdf GAL80* construct.

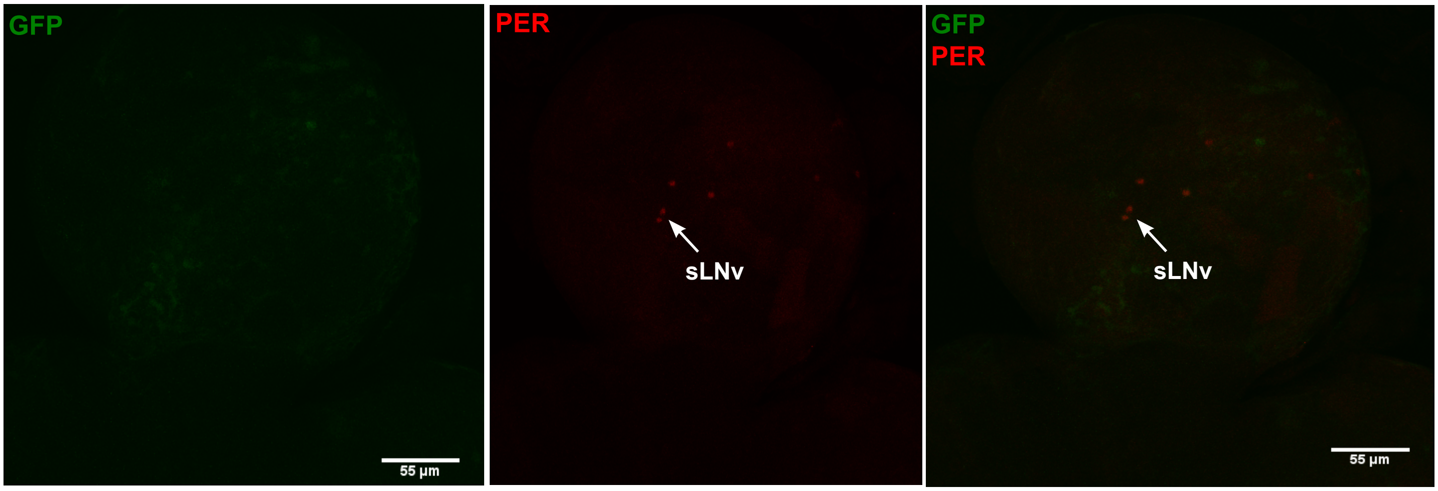


**Supplementary fig. S3:** Verification of the efficiency of *tub GAL80*$^{ts}$ construct

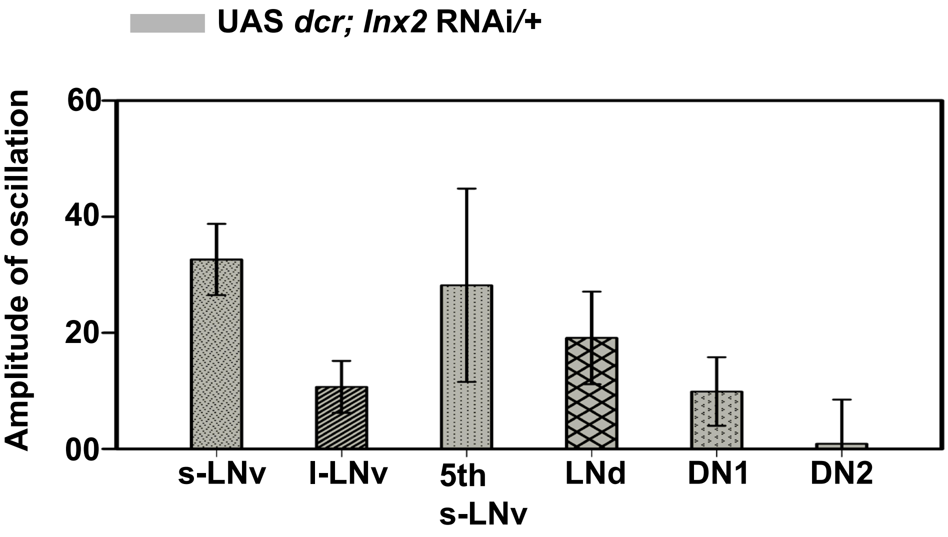


**Supplementary fig. S4:** Amplitude of PER oscillation in different circadian neuronal neuronal subsets in control flies on third day of DD.
